# Hand-hygiene mitigation strategies against global disease spreading through the air transportation network

**DOI:** 10.1101/530618

**Authors:** Christos Nicolaides, Demetris Avraam, Luis Cueto-Felgueroso, Marta C. Gonzalez, Ruben Juanes

## Abstract

Hand hygiene is considered as an efficient and cost-effective way to limit the spread of diseases and, as such, it is recommended by both the World Health Organization (WHO) and the Centres for Disease Control and Prevention (CDC). While the effect of hand washing on individual transmissibility of a disease has been studied through medical and public-health research, its potential as a mitigation strategy against a global pandemic has not been fully explored yet. In this study, we investigate contagion dynamics through the world air transportation network and analyze the impact of hand-hygiene behavioural changes of airport population against the spread of infectious diseases worldwide. Using a granular dataset of the world air transportation traffic, we build a detailed individual mobility model that controls for the correlated and recurrent nature of human travel and the waiting-time distributions of individuals at different locations. We perform a Monte-Carlo simulation study to assess the impact of different hand-washing mitigation strategies at the early stages of a global epidemic. From the simulation results we find that increasing the hand cleanliness homogeneously at all airports in the world can inhibit the impact of a potential pandemic by 24 to 69%. By quantifying and ranking the contribution of the different airports to the mitigation of an epidemic outbreak, we identify ten key airports at the core of a cost-optimal deployment of the hand-washing strategy: increasing the engagement rate at those locations alone could potentially reduce a world pandemic by 8 to 37%. This research provides evidence of the effectiveness of hand hygiene in airports on the global spread of infectious diseases, and has important implications for the way public-health policymakers may design new effective strategies to enhance hand hygiene in airports through behavioral changes.

## Introduction

In past centuries, contagious diseases would migrate slowly and rarely across continents. Black death, for example, which was the second recorded pandemic in history after the Justinian Plague, originated in China in 1334^1^ and it took almost 15 years to propagate from East Asia to Western Europe. While contagious diseases were then affecting more individuals within countries due to poor hygiene and underdeveloped medicine, the means of transportation of that era - sea and land - hindered the range and celerity of disease spreading. Nowadays in contrast, transportation means allow people to travel more often (either for business or for leisure) and to longer distances. In particular, the aviation industry has experienced a fast and continuing growth, permitting an expanding flow of air travelers. In 2017 alone, around 4.1 billion people traveled through airports worldwide^2^ while the International Air Transport Association (IATA) expects that the number of passengers will roughly double to 7.8 billion by 2036^3^. Transportation hubs such as airports are therefore playing a key role in the spread of transmittable diseases^4^. In severe cases, such disease-spreading episodes can cause global pandemics and international health and socio-economic crises. Recent examples of outbreaks show how quickly contagious diseases spread around the world through the air transportation network. Examples include the epidemic of SARS (Severe Acute Respiratory Syndrome) and the widespread H1N1 influenza. SARS initial outbreak occurred in February 2003, when a guest at a hotel in Hong Kong transmitted an infection to 16 other guests in a single day. The infected guests then transmitted the disease in Hong Kong, Toronto, Singapore and Vietnam during the next few days, and within weeks the disease became an epidemic affecting over 8,000 people in 26 countries across 5 continents^5, 6^. The H1N1 flu, which caused around 300,000 deaths worldwide^7^, had a similar timeline. The first confirmed case of H1N1 was reported in Veracruz, Mexico on April 2009, while within few days the infection migrated to the US and Europe, and two months later the World Health Organisation (WHO) and the Centers for Disease Control and Prevention (CDC) declared the disease as a global pandemic.

Bacteria, pathogens and viruses are also transmitted easily at airports or during flights, causing infections and bacterial diseases that can expand to global epidemics. The pathogens are transmitted through the air^8^, resulting in the contagion of airborne diseases, or through physical contact between individuals. The transmission is accelerated when a dense population of people is concentrated in a confined area^9^ like an airport, with lack of good hygiene and efficient air ventilation. After an outbreak, infections diffuse while infected individuals transmit the disease to susceptible individuals. Airports play a major role in such contagious dynamics^10^, as they contribute daily to the contact of people from all over the world, some of whom may be carrying endemic infections and bacteria from their country of origin. In addition, there are numerous highly contaminated surfaces which are frequently touched by the passengers at airports and inside aircrafts^11^. Self-service check-in screens, gate bench armrests, water fountain buttons and door handles at airports, as well as seats, tray tables and handles of lavatories in aircrafts, are all known to have high microbial contamination^12–14^.

Mitigation strategies are designed and implemented to inhibit a global pandemic. At the individual level there is a focus on behavioral change towards adopting different interventions in the case of a health emergency^15, 16^. Along with other developments in medicine, vaccination has made a big contribution in that direction leading to the extinction of past epidemics and a significant reduction of mortality due to specific infections^17^. Vaccination has a substantial mitigating effect when effective vaccines are available soon enough after the emergence of a new disease and when vaccination campaigns cover about 70% of a susceptible population^18^. However, despite the known impact of vaccines on the reduction of infections, the rate of vaccination in the population has remained unchanged over the past decade^19^. Social nudges such as peer effects or education on vaccination benefits, and changes in the design of vaccination campaigns, can be deployed to change human behavior towards the increase of influenza vaccination rates^20^. In addition of preventing disease spreading by vaccination, isolating patients at home or closure of high-risk places like schools can moderate the transmission of disease-causing pathogenic microorganisms.

Several actions within the world air transportation network can be implemented to control a disease spreading in the case of a health emergency^21^. At the global scale, mobility-driven interventions such as airport closures and deliberate rerouting of the travelers, can reduce the number of individuals passing through or traveling from/to regions where dangerous diseases prevail^22^. At the local scale, actions at individual airports such as frequent cleaning of public areas (e.g. toilets, gates, check-in desks, etc.), efficient air ventilation and enhanced sensitization of frequently touched surfaces can reduce the risk of contamination and the transmission of infections. Furthermore, personal hygiene is the most important factor to prevent the spread of an infection^23^. Coughing etiquette, face masks, no face touch and hand hygiene are the most common actions that air travelers can easily adopt. From those actions, hand washing is the simplest and most effective component for preventing the transmission of viruses^24–27^ and bacteria, and is regularly mentioned as the first recommendation of precautions in health care (see for example^28^). A scientific study on the effects of hand washing on the bacterial contamination of hands, showed that after a deliberate contamination of individuals by touching door handles and railings in public places, bacteria were found in 44% of the sample, while this percentage was reduced to 23% after hand washing with water only, and to 8% after hand washing with water and plain soap^29^. The same study concluded also that the effect of hand washing does not depend on the bacteria species.

While hand hygiene is considered as the first prevention step in the case of an epidemic emergency, there is lack of evidence for its effects as a mitigation strategy against global epidemic spreading. In this work, we study contagion dynamics through the world air transportation network and we elucidate the impact of hand-hygiene behavioral changes on the diffusion of infections worldwide. We develop a computational model to track the realistic mobility patterns of air travelers, their hand washing behavior and the dynamics of epidemic spreading. Human mobility is modelled by a stochastic agent-base system that accounts for the spatial distribution of airports, the realistic human mobility through the world air transportation network, and the waiting-time distributions of individuals at origin, destination and connecting airports. In addition, we develop a compartmental epidemic model which relates the disease spreading with hand-hygiene behaviour. Using Monte-Carlo simulations we assess the impact of hand washing at the early stages of a global epidemic spreading. From the simulation results, we measure the early-time spreading power of the 40 busiest airports under four different intervention scenarios: 1. increase of hand washing engagement homogeneously at all airports; 2. increase of hand washing engagement only at the source of the disease; 3. increase of hand washing engagement at the ten most important airports of the world air transportation network; and 4. increase of hand washing engagement at the ten most important airports for each source of the disease. The aim of this study is to identify the most effective mitigation strategy of hand hygiene contributing the most to the reduction of global epidemic spreading.

## Data description

We use world air traffic data provided by the Official Airline Guide (OAG), that includes all the trips (more than 1.9 million) that were booked in September 2017. Each row in the dataset states the number of passengers that traveled from an origin airport to a destination airport, and indicates any intermediate connecting flights (see Table 1 for example).

**Table 1.**
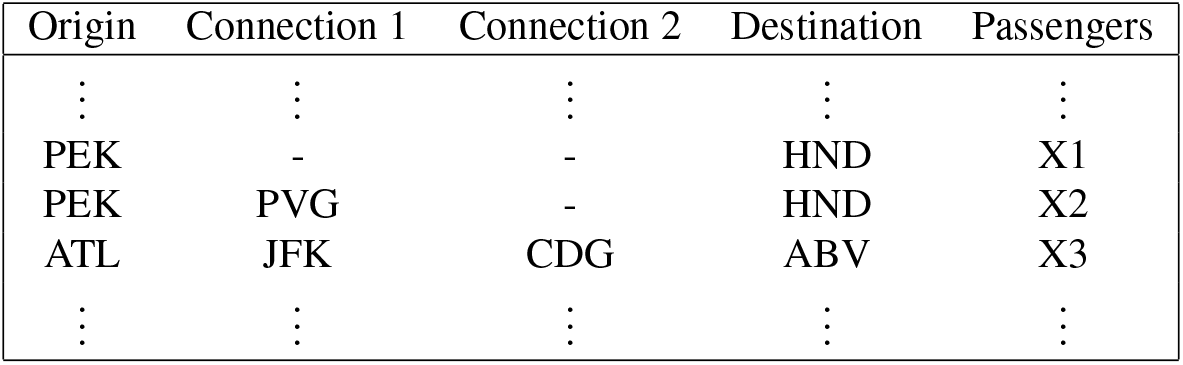
An example of three trips showing that in September 2017 X1 individuals traveled with direct flights from PEK (Beijing Capital International Airport, Beijing, China) to HND (Haneda Airport, Tokyo, Japan), X2 individuals traveled from PEK to HND with a layover at PVG (Shanghai Pudong International Airport, Shanghai, China), and X3 individuals traveled from ATL (Hartsfield-Jackson Atlanta International Airport, Atlanta, USA) to ABV (Nnamdi Azikiwe International Airport, Abuja, Nigeria) with connecting flights at JFK (John F. Kennedy International Airport, New York, USA) and CDG (Charles de Gaulle Airport, Paris, France) airports.

From the dataset, we observe that all trips in September 2017 were operated through a network of 3621 unique airports. For each airport, we estimate the total traffic by adding the number of passengers for the trips where the airport is denoted as ‘Origin’, the number of passengers for the trips where the airport is denoted as ‘Destination’, and twice the number of passengers for the trips where the airport is denoted as ‘Connection’ (either Connection 1 or 2). For subsequent computational efficiency, we restrict our analysis to the subset of the dataset corresponding to traffic among the 2500 *busiest* airports (by total traffic). This subset accounts for 98.25% of the total trips and 99.8% of the total traffic.

## Computational model

We build a computational model that simulates the mobility of travelers through the air transportation system, coupled with the propagation of a hypothetical infectious disease. Using the OAG data, we first generate the worldwide air transportation network, where the nodes are the 2500 busiest airports and the links between them are given by the connections between airports for which flights exist in the dataset. The network describes a heterogeneous metapopulation of airports where each individual airport is a subpopulation of individuals^30, 31^. We further develop a human mobility model to track the stochastic routes of traveling agents through the air transportation network. We finally implement a compartmental epidemic model to track the reaction dynamics of infection contagions as well as the hand washing related behavior of the traveling agents.

### Mobility Model

The human mobility model has the form of a stochastic agent-base tracking system^32, 33^ that accounts for the spatial distribution of airports, detailed air-traffic data, the correlated and recurrent nature of human mobility and the waiting-time distributions of individuals at different locations. We first generate the origin-destination *flux* matrix 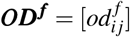 where 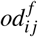 is the number of passengers that traveled in September 2017 from origin *i* to destination *j*, and the origin-destination *probability* matrix 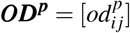 where 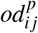 is the probability that an agent travels from origin *i* to destination *j*. Each element of the ***OD^p^*** matrix is calculated by 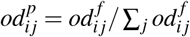 where 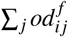 is the total number of passengers that traveled from origin *i*. We then assign a ‘home’ population *P_i_* at each subpopulation *i* following the nonlinear empirical relation 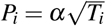, where *T_i_* is the total traffic at airport *i* and *α* is a constant parameter that is identified to give a total population size of *N* = Σ_*i*_*P_i_* individuals. In other words, each individual agent is initially assigned to its ‘home’ subpopulation i. Within the mobility route, the agent that was assigned to home *i* chooses to travel at a ‘destination’ airport *j* with probability extracted from the ***OD^p^*** matrix. If the two nodes *i* and *j* are connected by more than one path (i.e. direct when the two airports are connected with direct flights and indirect when the two airports are connected only with connecting flights), then the probability that the agent selects a given path is proportional to the relative number of passengers traveling in each direct or indirect flight from origin *i* to destination *j*. After each trip (from origin *i* to destination *j*), the agent returns back to its home airport. Thus, the stochastic mobility model generates the spatial trajectory for all agents. In addition, using realistic waiting times at the three distinct locations where an agent can be (i.e. home, connecting airport or destination) and actual flight times required to travel between the airports we express the spatio-temporal patterns of all the agents at the granularity of an hour. The waiting times at home airports, connecting airports and destinations are provided by the Bureau of Transportation Statistics 2010^34^, and follow right-skewed distributions with means 897.87 hours (~37 days), 1.33 hours, and 127.36 hours (~5 days) respectively. The average flight times between each airports *i* and *j*, are estimated as the ratio of the geographical distance of the two airports, *d_ij_*, calculated by the spherical law of cosines, over the average velocity of an airplane which is assumed to be constant and equal to 640 km/h considering the changes in takeoff, climb, cruise, descent and landing speeds.

### Epidemic Model

The conventional SIR model in epidemiology describes the reaction kinetics of an infection within a closed population^35^. According to the SIR model, each individual is considered as either susceptible (*S*), infected (*I*) or recovered (*R*). The sum of the compartments at any given time *t* is equal to the total population size (*S*(*t*) +*I*(*t*) + *R*(*t*) = *N*). The SIR reaction kinetics model two distinct processes: the infection process, 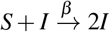, where an infected individual transmits the infection to a susceptible individual with rate β, and the recovery process, 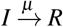, where an infected individual recover with rate *μ* (*μ*^−1^ is the average time required for an infected individual to recovers). The ratio *R*_0_ = *β*/*μ* defines the basic reproductive number of the infection, denoting the average number of secondary infections an infected individual causes before it recovers. For a closed subpopulation the disease dies out exponentially fast when *R*_0_ < 1, while in grows and potentially causes a pandemic for *R*_0_ > 1.

In this study, we modify the conventional SIR model to reflect the effects of hand washing behavior in the infection process. We formulate the SIR_WD_ model where each individual is placed in one of the three epidemic compartments (susceptible, infected, recovered) but also is characterized by one of the two hand cleanliness states, namely washed (*W*) or dirty (*D*) (Figure 1A).

**Figure 1.**
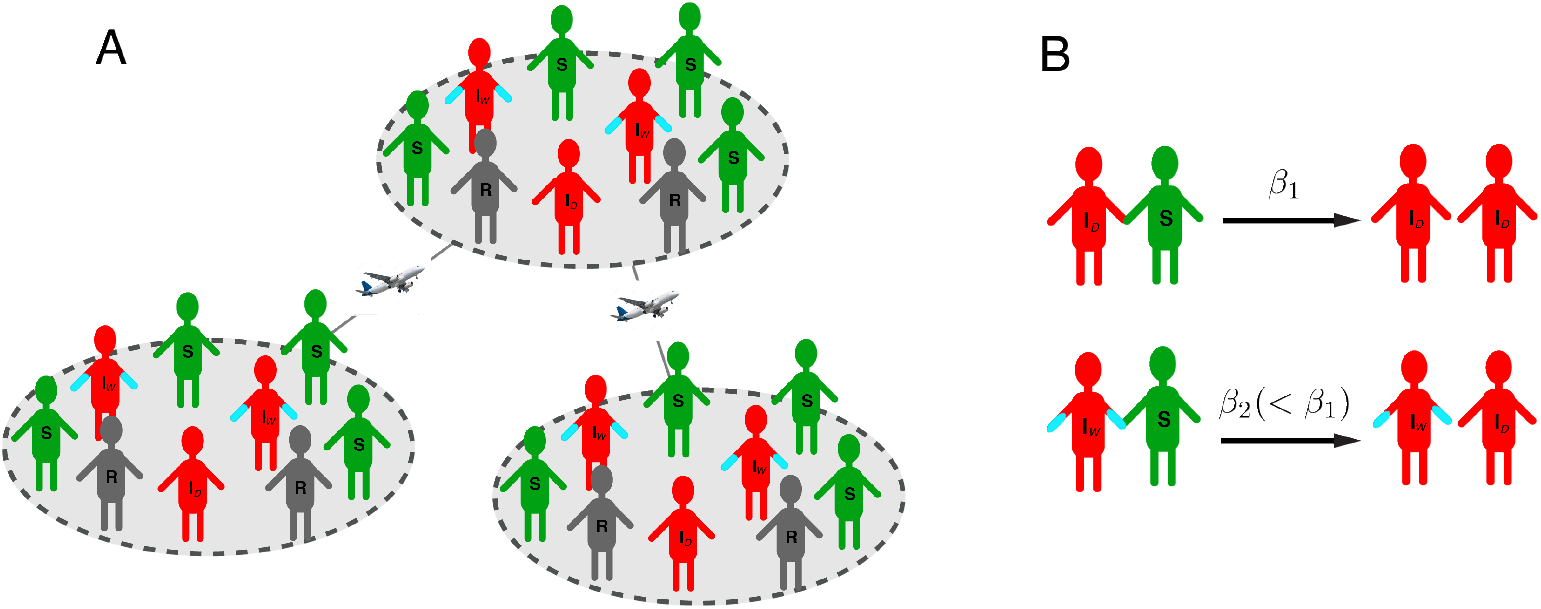
Pictorial demonstration of our model. (A) Illustration of the SIR_WD_ traveling population. Each individual can be either Susceptible to the disease, Recovered from the disease, Infected-Washed (blue hands) or Infected-Dirty. (B) Schematic diagram of the SIR_WD_ infection reaction. When an Infected-Washed individual comes in contact with a susceptible individual the probability of transmitting the disease is smaller compared to the case that has dirty hands.

The SIR_WD_ epidemic model is then expressed by:

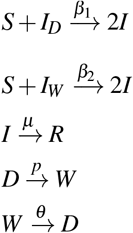

where *β*_1_ is the infection rate with which an infected individual with *dirty* hands transmits the infection to a susceptible individual (*β*_1_ is equal to the infection rate *β* of the conventional SIR model), *β*_2_ is the infection rate with which an infected individual with *washed* hands transmits the infection to a susceptible individual (*β*_2_ < *β*_1_), μ is the recovery rate (it is equal to the recovery rate of the conventional SIR model), *p* is the hand washing engagement rate (denoting the percentage of individuals with non-clean hands that move to the washed state within the next hour) and θ is the hand washing effectiveness rate (*θ*^−1^ denotes the average duration that an individual with washed hands returns back to the ‘dirty’ state). The infection reactions that are described in the first two expressions of the SIR_WD_ model are shown in the diagram of Figure 1B.

To get the infection, a healthy individual needs to touch to a contaminated surface or directly an infected person. If the individual is healthy and touch a contaminated surface — independent of how long ago he/she washed his/her hands — he/she will get the bacteria on hands. However, if he/she washes hands soon after he/she gets contaminated there is big probability of taking that bacteria out of hands before they are transmitted to body fluids. Therefore, the hand washing rate of healthy individuals affects the transmissibility of a disease as well. Our SIR_WD_ model takes into account only the interdependence between disease transmission probability and the hand cleanliness of the infected individuals. To model the process where the hand washing behavior of susceptible/healthy individuals has a role in the infection process, we need to build a more sophisticated model based on SEIR reaction kinetics, where the extra epidemic compartment *E* indicates individuals that are *Exposed* to bacteria or viruses^36^. The SEIR epidemic model describes the following three processes: (a) a susceptible comes in contact with an infected individual and becomes exposed to the disease with some rate 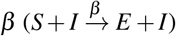, (b) an exposed becomes infected with some rate 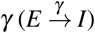, and (c) an infected recovers with rate 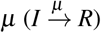. Both rates *β* and *γ* are affected by the hand washing levels. Here, we keep our analysis simple by using the conventional SIR model with the assumption that if infected individuals wash hands frequently, there is smaller probability to contaminate surfaces or other healthy people directly.

### Initial conditions and assumptions

We assume a flu-type disease, where the recovery rate is *μ* = 1/4 days^−1^ (i.e. on average each infected individual recovers after four days) and the reproductive number is *R*_0_ = 3 (i.e. on average each infected individual transmits the disease to three other individuals). The infection rate for the SIR model is *β* = *μR*_0_ which is equal to the infection rate *β*_1_ for the processes 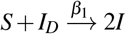 in the SIR_WD_ model. The infection rate of the process 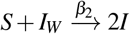 is *β*_2_ = 0.6*β*_1_ as effective hand washing has been proven to be able to prevent around 50-70% of infections^37^. The hand washing effectiveness rate which indicates the average time that washed hands become again contaminated is set to *θ* = 1/1.5 hours^−1^ (i.e. is the rate to change from ‘washed’ to ‘dirty’ state). We also consider that mostly 1 over 5 people in an airport have cleaned hands at any given moment in time (i.e. 20% of airport population). This is equivalent to hand washing engagement rate among the non-cleaned individuals equal to *p* = 0.12 per hour (i.e. every hour about 12% of the non-cleaned individuals are washing their hands). We declare this hand washing engagement rate (*p* = 0.12 hours^−1^) as the *status quo* (see next section). We vary *p* to analyze and quantify the effect of hand washing engagement on different scenarios of epidemic spreading.

### Status quo of hand washing engagement rate

To derive an approximation of the status quo level of hand cleanliness (i.e. the percentage of people with cleaned hands) in the population of an airport at any given moment, we simulate a close population following some assumptions derived from the literature. We use data from a survey performed by the American Society for Microbiology^38^ which revealed that 30% of travelers do not wash their hands after using public toilets at airports, denoting that the rest 70% are compliers with hand washing. Following a study in a college town environment, we consider that only the 67% of the compliers wash their hands properly (i.e. with water and soap and for the recommended by CDC duration of time^39^), while the rest 33% are wetting their hands quickly and/or without soap^40^. Therefore, we assume that in an airport population of *N* individuals only the 70% · 67% = 49.6% of *N* are compliers with effective hand washing. Furthermore, we assume that each individual wash their hands on average between 4-10 times per day^41^ which means that in a 24-hour timeframe one event of hand washing takes place every 2.5-6 hours. We assume that the frequency of hand washing follows a normal distribution with mean equal to 4.5 hours and standard deviation equal to 1 hour. We also consider that the duration of cleanliness of hands after hand washing follows an exponential distribution with mean value equal to 1.5 hours.

Using the above approximations, we find that that at any given moment, the percentage of passengers in an airport that have cleaned hands has an *upper bound* of 24%. Given that this is very optimistic upper bound of the reality, we assume and use in simulations that the status quo for the percentage of individuals that have clean hands in an airport at any given moment is 20%. To preserve a stable 20% hand cleanliness level over time in an airport, the *hand washing engagement rate* in the compartmental SIRWD model, that indicates the rate of hand washing per hour between individuals with non-cleaned hands, is calculated to be equal to *p* = 0.12 h^−1^ (i.e. 12% of dirty individuals wash their hands within an hour). This indicates the status quo of hand washing engagement rate. In the case that we would like to increase the level of hand cleanliness in an airport to 30% or 40% or 50% or 60% we need to increase the hand washing engagement rate to 0.21 h^−1^ or 0.32 h^−1^ or 0.49 h^−1^ or 0.73 h^−1^ respectively.

### Methodology

We implement the epidemic model within the mobility model using Monte-Carlo simulations to track the mobility and contagion dynamics through the air transportation network. In the simulations, we consider different hand-hygiene mitigation strategies and we study their effects on the propagation and the diffusion of a disease at the global scale. We first study the conventional SIR epidemic model to identify the spatio-temporal structure of the disease for different seeding scenarios and to identify the most influential spreaders within the air transportation network. Furthermore, we study four hand-hygiene scenarios and their effectiveness to disease spreading inhibition: a. homogeneous increase of hand washing engagement at all airports, b. increased hand washing engagement at the ten most influential airports in the network, c. increased hand washing engagement at the ten most influential airports for each source of the disease, and d. increased hand washing engagement only at the source of the disease.

### Monte-Carlo simulations

At the initial time step of each simulation, *t* = 0, we declare an airport *i* as the source of the disease where we randomly choose ten individuals to seed the infection. For each analysis we run 500 realizations of 10^5^ traveling agents each. At each time step, which corresponds to *one hour*, we let individuals travel, wash their hands, and recover or transmit the disease to susceptible agents when those individuals are infected. At each time step, an infected individual recovers with probability Π_*I*→*R*_ = 1 − exp(−*μ*). When the transmission of an infection is associated with hand cleanliness of the infected individuals (as described by the SIRWD model), the probability of a susceptible to get the infection is Π_*S*→*I*_ = (1 − (1 − *β*_1_/*N_i_*)^*I_D,i_*^) + (1 − (1 − *β*_2_/*N_i_*)^*I_W,i_*^, where *I_D,i_* and *I_W,i_* are the numbers of ‘dirty’ and ‘washed’ infected individuals respectively at airport *i* and *N_i_* is the total population at airport *i*. The probability that an individual with washed hands becomes dirty is Π_*W→D*_ = 1 − exp(−*θ*) and the probability that an individual with dirty hands will wash his/her hands, within each one-hour time step, is Π_*D→W*_ = 1 − exp(−*p*). Using these probabilities, the computational model generates the stochastic epidemic transitions for the traveling agents over time. In our analysis, we vary the model parameter *p*, considering different hand-hygiene interventions, and analyze their impact on global disease spreading.

### Evaluating the early-time impact of the disease

We evaluate the early-time impact of the disease by measuring two quantities that are correlated between them: the *disease prevalence* and the *Total Square Displacement* two weeks after the disease is deliberately seeded in a source. The disease prevalence (PREV) is given by the total number of affected individuals (infected plus recovered)^42^. However, as we want to evaluate not only the total number of infected individuals but also how well spread they are within the globe, we use the Total Square Displacement (TSD) of the infected individuals as a simulation metric^32^. This metric is given by the formula 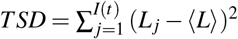, where *I*(*t*) is the number of infected individuals at time *t* = 2 weeks, *L_j_* is the geographic location of the *j*-th infected individual and 〈*L*〉 is the position of the geographic centre of the infection. The geographic centre is the centre of gravity (aka the centre of mass) for the locations of all infected individuals. To find the geographic centre, we first convert the latitude and longitude of each location *Lj* from degrees to radians, and then into Cartesian coordinates using the formulas *x_L_j__* = cos(*lat_L_j__π*/180) · cos(*lon_L_j__π*/180), *y_L_j__* = cos(*lat_L_j__π*/180) · sin(*lon_L_j__π*/180) and *z_L_j__* = sin(*lat_L_j__π*/180). We then calculate the mean of the Cartesian coordinates by 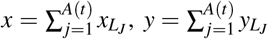 and 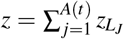, and finally we convert the average coordinates (*x, y, z*) into latitude and longitude in radians using the four-quadrant inverse tangent function 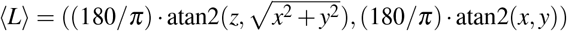.

## Results

### Conventional SIR Model

The contagion dynamics of infectious diseases are broadly described by the basic SIR model. However, the concept of SIR reactions excludes the effects of individual hygiene activities (like hand washing) from the model of infection transmissions. In that case, the infection reaction process is considered as independent from the hand cleanliness of the infected individuals. In our initial analysis, we first use the SIR model (disregarding the impact of hand-hygiene behavior) to estimate the capacity of airports to spread an infectious disease globally. We seed the disease in each of the world major airports and through simulations we track the contagion dynamics two weeks after the outbreak. We rank the airports according to their spreading capacity as quantified by the TSD of infected individuals^32^ (Figure 2 – middle). From the analysis, it is observed that the total traffic alone cannot predict the power of an airport to spread the disease (comparing left and middle panels in Figure 2), but should be accounted alongside with the location of each spreader airport and the spatial correlations with other influential airports in the network. NRT (Narita International Airport, Tokyo, Japan) and HNL (Honolulu International Airport, Honolulu, USA) airports are indicative examples, as they ranked in the 46th and 117th place by total traffic respectively, but they have large contribution in the acceleration and expansion of a disease contagion globally (ranked by TSD on the 7th and 30th place respectively). This happens because NRT and HNL combine three important features with high impact on the disease spreading. These are (i) they have direct connections with the world’s biggest mega-hub airports, (ii) they operate long-range in- and out-bound international flights, and (iii) they are located in geographical conjunctive points between East and West^32^.

**Figure 2.**
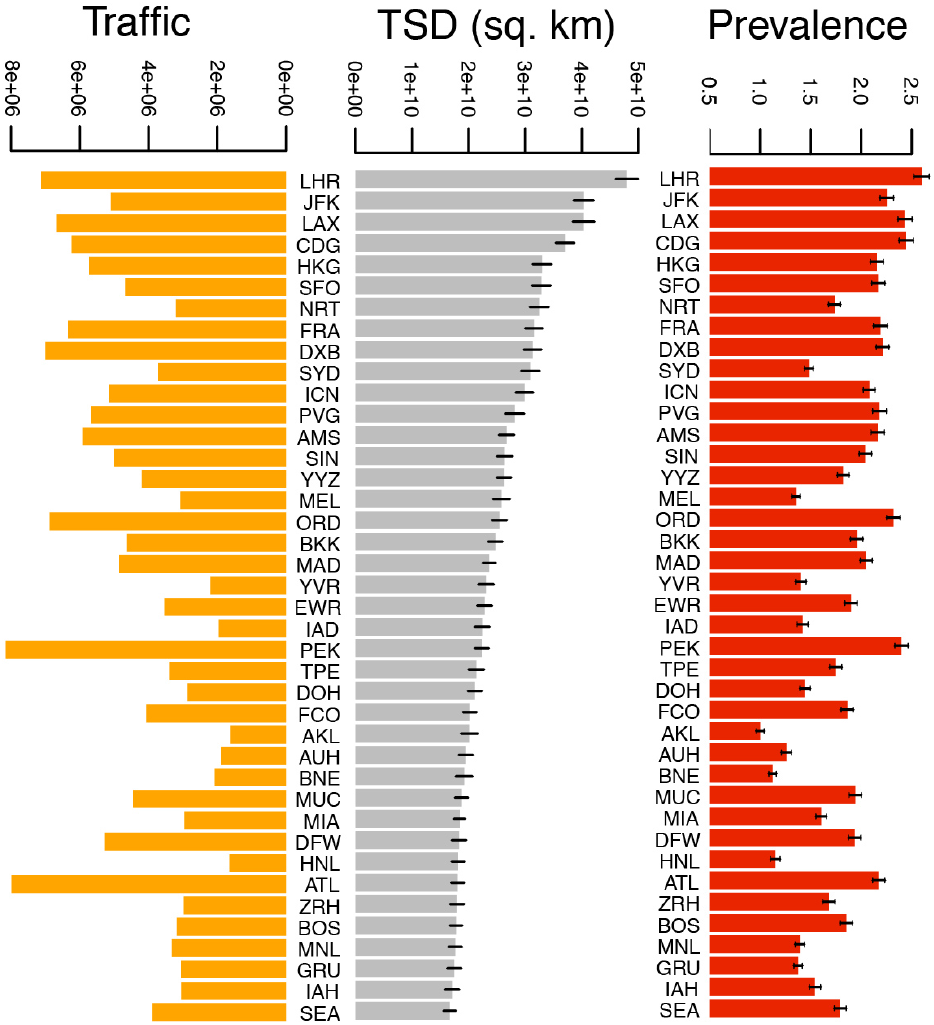
The impact of the source of the disease on its global spread. (middle) Ranking of the 40 most influential airports in the world with respect to the TSD of the infected individuals two weeks after the disease started from each one of these airports. (right) The two-week prevalence of the disease as measured by the percentage of world population that have been affected (infected plus recovered) by the disease two weeks after a disease started from each of these major airports. (left) The total monthly traffic of each of those major airports as has been calculated using the world air-traffic dataset from September 2017.

The bar plot in the right of Figure 2, shows the two-week prevalence of the disease as measured by the percentage of world population that have been affected by the disease two weeks after a disease started from each of the major airports. The two-week prevalence is highly correlated with the total traffic of the airport (the Pearson correlation coefficient is equal to 0.88) indicating that large airports have a big impact in terms of absolute number of affected (infected plus recovered) individuals.

### SIR_WD_ Model: Worldwide homogeneous hand washing intervention

The effects of hand-hygiene are then embedded in the computations and we focus the analysis on the epidemic reaction kinetics as described by the SIR_WD_ model. For each simulation, the disease is seeded in one of the major airports (ten randomly chosen individuals are infected *t* = 0) and the epidemic expansion all over the world due to the mobility of infected agents is recorded. In the status quo scenario, we consider that hand cleanliness level is on a 20% steady state at each airport in the world. This percentage represents the fraction of individuals with washed hands at any moment. The rate of hand washing per hour that corresponds to 20% cleanliness is equal to 0.12 h^−1^ (see Table 2). We rank the airports in respect to TSD metric and we observe that LHR has the greatest impact while LAX, JFK, SYD and CDG are among the five most influential spreaders worldwide. Using the same order of airports, we repeat the simulations, by increasing the hand washing engagement rate homogeneously at all airports to achieve global hand cleanliness levels of 30%, 40%, 50% and 60%. For each hand washing engagement rate (or hand cleanliness level) we analyse the changes in the impact of contagion.

Figure 3A shows the early-time evolution of the fraction of affected individuals over the first two weeks after a disease is seeded at DXB (Dubai Airport). Its observed that with the increase of hand cleanliness level at all airports from 20% to 60% there is a significant reduction in the percentage of the affected individual in the total population from around 1.5% to less than 0.5%. In Figure 3B we demonstrate the spreading power of the most influential spreader airports measured by TSD of infected individuals two weeks after a disease was initiated at each of these major airports, at 20% (status quo), 30%, 40%, 50% and 60% of homogeneous hand cleanliness. A drastic, very significant reduction in TSD is observed with the increase of cleanliness level, verifying that hand-hygiene is one of the most important factors to control or even prevent an infection. For example, the spread of infection seeded in LHR was covered about 5 · 10^5^ square meters around the mass centre of the infection within two weeks, while infected area was reduced to less than 2 · 10^5^ square meters when cleanliness level increased from 20% to 60% globally. The relative to the status quo reduction of the disease impact is calculated by (*TSD*_20%_ − *TSD_X_*)/*TSD*_20%_ for the TSD metric or (*PREV*_20%_ − *PREV_X_*)/*PREV*_20%_ for the disease prevalence metric, where the cleanliness level *X* increases from 30% to 60% worldwide. The results are shown in Table 2 indicate a significant reduction of the impact of a disease worldwide by 24% to 69% depending on the hand washing engagement rate worldwide using the calculated by the TSD (or by 18% to 55% as calculated by the global prevalence of the disease).

**Figure 3.**
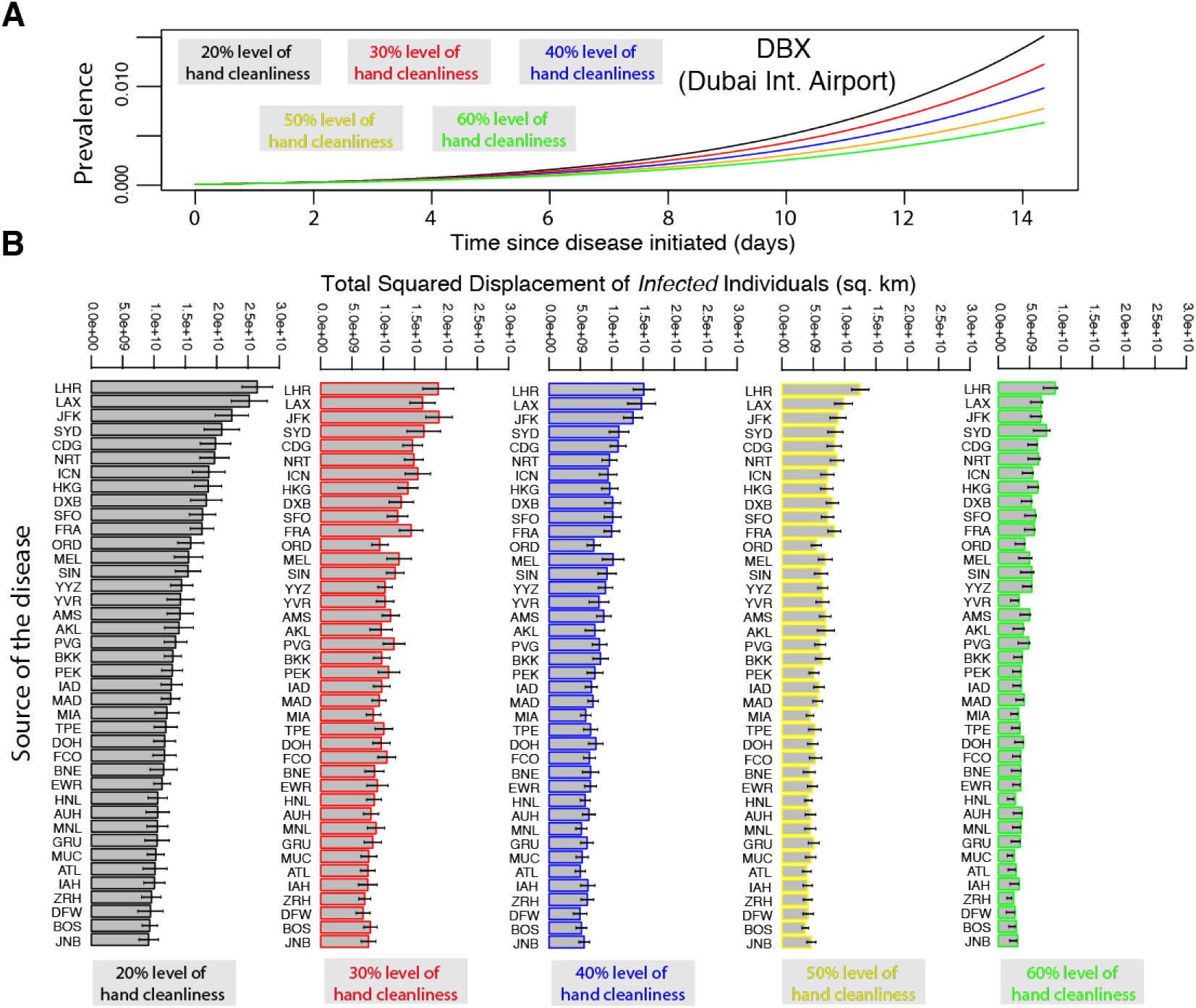
The effect of a global, homogeneous hand washing strategy on the impact of a disease spreading. (A) The fraction of affected (infected plus recovered) individuals worldwide over the first two weeks after the infection was initiated at Dubai International Airport at different levels of hand cleanliness. (B) Airports are ranked according to their spreading power to transmit a disease faster and further across globe – measured by the total squared displacement of infected individuals two weeks after a disease started from each individual airport. From left to right the hand cleanliness level increases from 20% (status quo) to 60%.

**Table 2.**
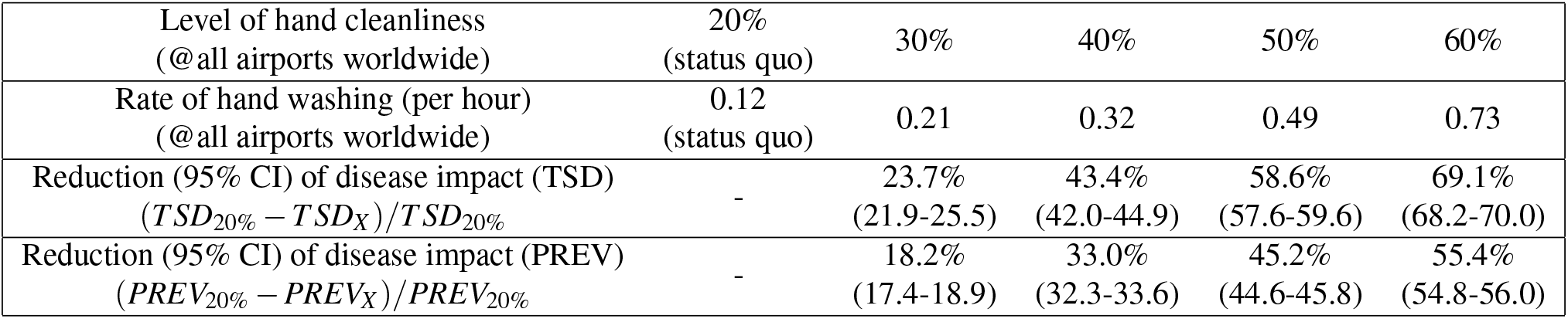
Reduction of the disease impact with a homogeneous increase of hand washing engagement worldwide. These are point estimates and 95% Confidence Intervals calculated across 120 disease spreading scenarios. In each scenario, the source of the disease is one of the 120 largest airports in the world. Each spreading scenario is evaluated over 100 mobility and epidemic realizations.

|  |  |  |  |  |  |
| --- | --- | --- | --- | --- | --- |
| Level of hand cleanliness<br>(@all airports worldwide) | 20%<br>(status quo) | 30% | 40% | 50% | 60% |
| Rate of hand washing (per hour)<br>(@all airports worldwide) | 0.12<br>(status quo) | 0.21 | 0.32 | 0.49 | 0.73 |
| Reduction (95% CI) of disease impact (TSD)<br>$(TSD_{20\%} - TSD_X)/TSD_{20\%}$ | - | 23.7%<br>(21.9-25.5) | 43.4%<br>(42.0-44.9) | 58.6%<br>(57.6-59.6) | 69.1%<br>(68.2-70.0) |
| Reduction (95% CI) of disease impact (PREV)<br>$(PREV_{20\%} - PREV_X)/PREV_{20\%}$ | - | 18.2%<br>(17.4-18.9) | 33.0%<br>(32.3-33.6) | 45.2%<br>(44.6-45.8) | 55.4%<br>(54.8-56.0) |

### SIR_WD_ Model: Strategic hand washing policies

While increasing the level of hand washing engagement homogeneously at all airports is very costly and maybe infeasible, we test some other less costly intervention strategies. These interventions consider the increase of hand washing engagement rate only at a small number of ‘key’ airports. We test three different intervention strategies that consider the increase of the hand washing engagement rate: (i) at the ten pre-identified key airports worldwide, (ii) the ten key airports of each source of the disease, and (iii) only at the source of the disease.

For intervention scenario (i), we pre-identify the ten key airports of the world air transportation network by multiplying the susceptibility of each airport by the strength of the airport to spread an infection globally. The strength of airport *i* is calculated as 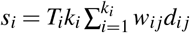, where *T_i_* is the total outgoing traffic from airport *i*, *k_i_* is the number of connections of *i* (i.e. the degree of node *i* in the network), and 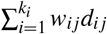 is the effective length of all links of *i* which is the weighted sum of the actual distances *d_ij_* between *i* and *j* nodes. The weights *w_ij_* are the fractions of passengers traveling from *i* to j. The susceptibility of airport *i* is calculated using the conventional SIR simulations as the weighted average fraction of infected individuals that arrive at *i* over all the seeding scenarios considered in the SIR model described above. Using the above combined metric (susceptibility × strength), we identify the ten ‘key’ airports of the world air transportation network as being the LHR, LAX, JFK, CDG, DXB, FRA, HKG, PEK, SFO and AMS. For the intervention scenario (ii), we identify ten ‘key’ airports for each source of the disease, by multiplying the airport strength by the source-dependent susceptibility. The source-dependent susceptibility of airport *i* for the seeding of the disease at airport *j* is calculated as the fraction of infected individuals that arrive at *i* when the disease is initiated at airport *j*. Therefore, for this intervention scenario, knowledge of the source of the disease is required and for different sources of the disease we have different sets of ‘key’ airports (see Figure 4). Finally, for the intervention scenario (iii), since we increase the hand washing engagement rate only at the source of the disease, prior knowledge of the source is required.

**Figure 4.**
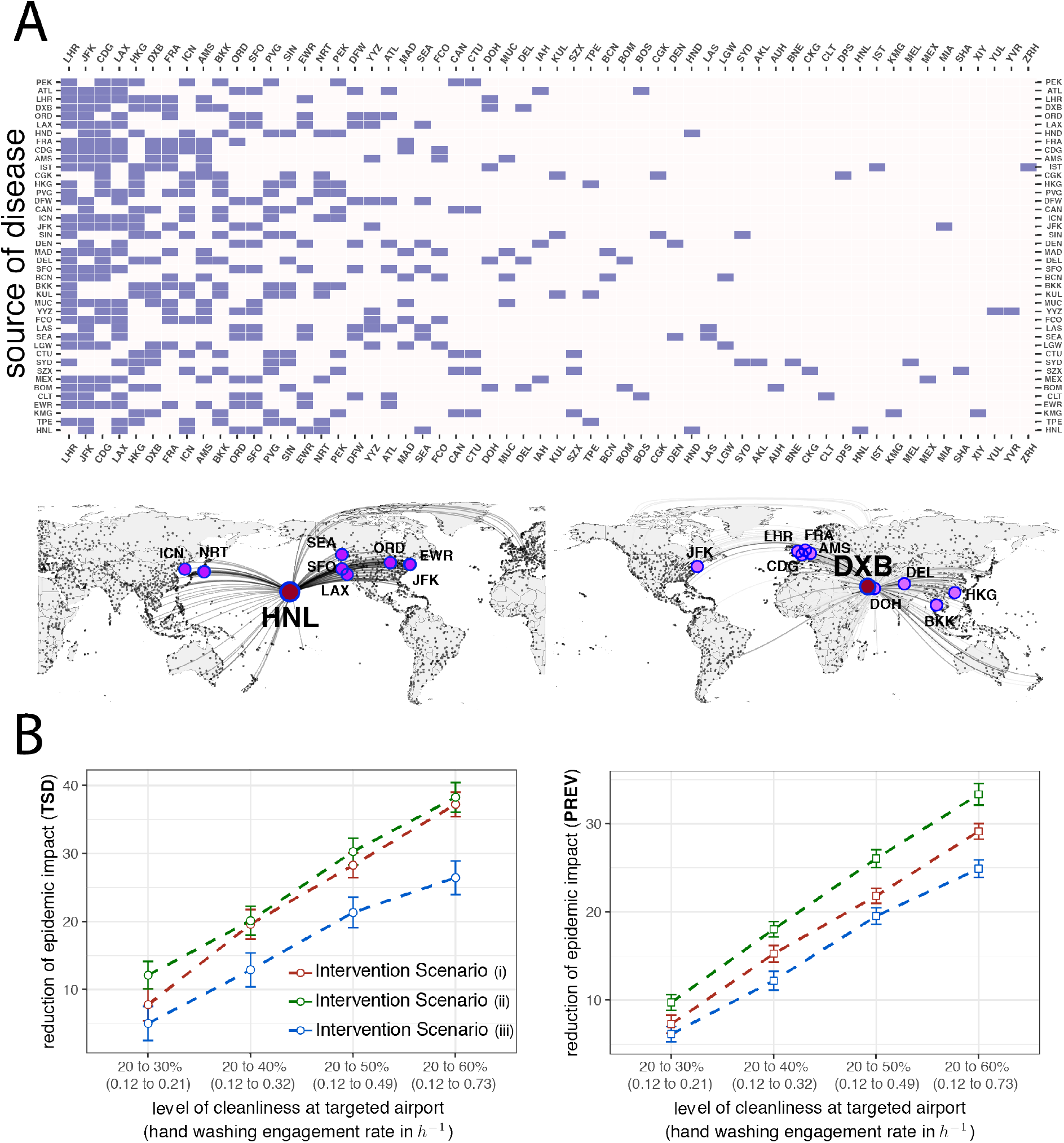
The effect of strategic hand washing policies on the impact of a disease spreading. (A) The ten key airports of each source of disease. When the disease is seeded in each of the source (in this plot we show as source the 42 busiest airports of the network), we increase the hand washing engagement rate at the ten key airports in relation to each source for scenario (ii) of our simulations and analyse the early-time contagious dynamics. (lower) The locations of the ten important airports for HNL - Honolulu International Airport (left) and for DXB - Dubai International Airport (right) shown in the global map. (B) Reduction of the disease impact as a function of the level of hand cleanliness (or hand washing engagement rate) with respect to status quo for the three different intervention strategies (scenarios). Disease impact is calculated with respect to the Total Square Displacement (TSD) at the left and the Prevalence of the disease (PREV) at the right. These are point estimates and 95% Confidence Intervals across 120 disease spreading scenarios. In each spreading scenario, the source of the disease is one of the 120 largest airports in the world. Each seeding scenario is evaluated over 100 mobility and epidemic realizations.

The results, shown in Figure 4, indicate that the design of a less costly (compared to homogeneous) strategic plan for hand washing intervention only at ten pre-identified “key” airports worldwide (Scenario (i)) could lead to a significant reduction of the disease impact calculated by the TSD from ~8% to ~37% (or ~7% to ~29% calculated by the world prevalence). If the strategic plan is deliberately implemented only at the ten most important airports for each source of disease (Scenario (ii)), we observe a further reduction of the disease impact. However, this further reduction is statistically different from that of Scenario (i) only in terms of the Prevalence of the disease, and not in terms of geographical spreading as calculated through TSD. Intervention Scenario (iii), that considers enhancing hand washing engagement only at the source of the disease also has a significant effect on the reduction of disease impact; yet, this effect is smaller than that of intervention Scenarios (i) and (ii).

## Discussion

In this work we have analysed contagion dynamics through the world air transportation network and the impact of hand-hygiene behavioural changes of air travelers against global epidemic spreading. Using well-established methodologies, we have applied simulations to track traveling agents and their hand washing activity and analysed the expansion of flu-type epidemics through the world air transportation network. From the simulation results, we have measured the early-time spreading power of the major airports in the world under different hand-hygiene interventions. Using data-driven calculations, we estimated that mostly 1 over 5 people are cleaned at any given moment in time (i.e. 20% of airport population). This is translated to hand washing engagement rate among the non-cleaned individuals equal to 0.12 per hour (i.e. every hour about 12% of the non-cleaned individuals are washing their hands). From simulation results we have shown that, if we are able to increase the level of hand cleanliness at all airports in the world from 20% to 30% (or equivalently to increase the hand washing engagement rate from 0.12 to 0.21 per hour), either by increasing the capacity of hand washing and/or by increasing the awareness among individuals and/or by giving the right incentives to individuals, a potential infectious disease will have a worldwide impact that is about 21.2% smaller compared to the impact that the same disease would have with the 20% level of hand cleanliness (or 0.12 per hour hand washing engagement rate). Increasing the level of hand cleanliness to 60% (or equivalently the hand washing engagement rate among non-cleaned individuals to 0.73 per hour) at all airports in the world would have a reduction of 64.6% in the impact of a potential disease spreading. Moreover, we have identified the ten most important airports of the network, for which increasing the level of hand cleanliness (or hand washing engagement rate) only at those, the impact of the disease spreading would decrease by 9% to 37%.

Our current analysis has some limitations, which can be addressed in the future. A first limitation is the use of the simple SIR reaction kinetics while a more complicated model, like the SEIR, will provide inferences on the impact of hand washing behavior among the individuals exposed to the disease on the expansion of epidemics. A second limitation is that we use data from the air transportation system as a proxy for human mobility. A complete analysis should focus on the spread of infections through a more realistic human mobility network that includes daily commuting patterns and travel through other means of transportation. A third limitation is the assumption of a homogeneous hand-hygiene behavior of air travelers, as we do not know the actual hand washing activity that varies between individuals within a local population and between individuals from different societies and cultures. Future research can be designed to understand the human hand-washing behavior and provide insights on what and how social effects can change it.

Epidemiological outbreaks not only increase global mortality rates, but they also have a large socio-economic impact that is not limited to those countries that are directly affected by the epidemic. Outbreaks reduce the consumption of goods and services, negatively affecting the tourism industry, increasing businesses’ operating costs, and speeding the flight of foreign capital, generating massive economic costs globally. For instance, even the relatively short-lived SARS epidemic in 2003 led to the cancellation of numerous flights and to the closure of schools, wreaking havoc in Asian financial markets and ultimately costing the world economy more than $30 billion^43^. Hypothetical scenarios of future global pandemics give estimates on the economic effects. The worldwide spread of a severe infectious disease is estimated to cause approximately 720,000 deaths per year and an annual reduction of economic outcome of $500 billion (i.e. ~0.6% of the global income)^44^. In such severe scenarios where markets shut down entirely, a massive global economic slowdown is expected to occur shrinking the GDP of national economies. Of course, wealth and income effects are expected to differ sharply across countries, with a major shift of global capital from the affected economies (i.e. of developing countries) to the less-affected economies (i.e. of North America and Europe).

The effectiveness of mitigation strategies against global pandemics is evaluated through the total expected cost versus the total public health benefit^45^. The target of each strategy is to maximise the social welfare by incurring in the minimum economic cost. For interventions where travel restrictions are implemented^46^, the cost increases with the number of closed airports and the number of individuals that get stranded in those airports. The reward is related to the relative decrease in the global footprint of the disease, compared with the null case of non-interventions. In contrast to the mobility-driven strategies that change the population’s mobility patterns, other solutions such as hand washing appear to be more cost- and reward-effective. A future research on the socio-economic impact of global pandemics and the cost-effectiveness ratio of different mitigation strategies (e.g. hand washing, vaccination, airport closures, mobility routing diversions) against disease spreading would evaluate the efficiency and significance of hand-hygiene interventions. However, while hand hygiene is considered as the first prevention step in the case of an epidemic emergency, the capacity of hand washing facilities in crowded places, including airports, is limited only to wash basins at restrooms. It is not known, however, if increased capacity would enhance hand washing engagement by air travelers. New technology is being developed aiming to increase the capacity of facilities even outside restrooms, thus expanding the options for hand hygiene and the solutions for air and surface sterilization. Airbus^47^, for example, is exploring an innovative antimicrobial technology that is able to eliminate viruses and pathogens from aircraft surfaces (e.g. tray tables, seat covers, touch screens, galley areas). Boeing is also exploring a prototype self-sanitizing lavatory that uses ultraviolet light to kill 99.99% of pathogens^48^. At the same time, robotic systems for dirt detection and autonomous cleaning of contaminated surfaces^49^ and smart touch-free hand washing systems^50^ are promising tools on the evolution of cleaning technologies.

An important question is how such smart technologies are adopted by the general public, and what incentives can promote hand washing behavioral changes. Do digital nudges (motivation messages) make health related establishments attractive to individuals? A recent study has found that nudges have been effective at improving outcomes in a variety of health-care settings including a significant increase of influenza vaccination rates^20^. Can social influence or peer effects improve smart hand-washing engagement? Recent works have identified that social influence plays an important role in many behaviors like exercise or diet^51, 52^, and there is some initial evidence that it can play a role on individual hygiene^53^. There is certainly a need for rigorous and carefully designed field experiments on large population scale to identify and measure the causal effect of digital nudges, incentives and peer influence on public hand washing engagement of air travelers as well as the mechanisms of health-enhancing human behavior change.

This research can potentially shape the way policymakers design and implement strategic interventions based on promoting hand washing in airports that will lead to hindering any infection within a confined geographical area at the early days of an outbreak and inhibit the expansion as a pandemic. The most important outcome derived from our study is the conclusion that proper hand-hygiene with regular and efficient hand washing is the simplest and most effective solution for preventing transmission of infections and reducing the chances of massive epidemics spreading globally. This should be followed up by the design of strategic mechanisms able to increase the capacity of hand washing facilities in public places, and nudges that will enhance the adoption of hand-hygiene related behaviors.

## Acknowledgements

C.N. and D.A. acknowledge funding by SMIXIN Inc, a clean-tech startup that provides smart hand washing solutions. L.C.-F. and R.J. gratefully acknowledge funding from the MIT International Science and Technology Initiatives, through a Seed Fund grant. The authors would also like to acknowledge OAG (https://www.oag.com/) for providing the air-traffic data. The authors declare no competing interests.

## References

1. Centers for Disease Control and Prevention. History of the plague. https://www.cdc.gov/plague/history/index.html (2015). Online; accessed October 2018.

2. International Civil Aviation Organization. Continued passenger traffic growth and robust air cargo demand in 2017. https://www.icao.int/Newsroom/Pages/Continued-passenger-traffic-growth-and-robust-air-cargo-demand-in-2017.aspx (2018). Online; accessed October 2018.

3. International Air Transport Association. 2036 forecast reveals air passengers will nearly double to 7.8 billion. https://www.iata.org/pressroom/pr/Pages/2017-10-24-01.aspx (2017). Online Press Release No 55; accessed October 2018.

4. Brownstein, J. S., Wolfe, C. J. & Mandl, K. D. Empirical evidence for the effect of airline travel on inter-regional influenza spread in the United States. PLoS Medicine 3, e401, DOI: https://journals.plos.org/plosmedicine/article?id=10.1371/journal.pmed.0030401 (2006).

5. Peiris, J. S. M., Guan, Y. & Yuen, K. Y. Severe acute respiratory syndrome. Nat. Medicine 10, S88–S97, DOI: http://dx.doi.org/10.1038/nm1143 (2004).

6. World Health Organisation. SARS (Severe Acute Respiratory Syndrome). http://www.who.int/ith/diseases/sars/en/. Online; accessed October 2018.

7. Dawood, F. S. et al. Estimated global mortality associated with the first 12 months of 2009 pandemic influenza A H1N1 virus circulation: a modelling study. The Lancet Infect. Dis. 12, 687–695, DOI: 10.1016/S1473-3099(12)70121-4 (2012).

8. Memish, Z. A. et al. Environmental sampling for respiratory pathogens in Jeddah airport during the 2013 Hajj season. Am. J. Infect. Control. 42, 1266–1269, DOI: https://doi.org/10.1016/j.ajic.2014.07.027 (2014).

9. Dalziel, B. D. et al. Urbanization and humidity shape the intensity of influenza epidemics in U.S. cities. Science 362, 75–79, DOI: 10.1126/science.aat6030 (2018).

10. Lawyer, G. Measuring the potential of individual airports for pandemic spread over the world airline network. BMC Infect. Dis. 16, 70, DOI: 10.1186/s12879-016-1350-4 (2016).

11. Ikonen, N. et al. Deposition of respiratory virus pathogens on frequently touched surfaces at airports. BMC Infect. Dis. 18, 437, DOI: 10.1186/s12879-018-3150-5 (2018).

12. Zhao, B. et al. Microorganisms @ materials surfaces in aircraft: Potential risks for public health? – A systematic review. Travel. Medicine Infect. Dis. DOI: https://doi.org/10.1016/j.tmaid.2018.07.011 (2018).

13. McKernan, L. T., Burge, H., Wallingford, K. M., Hein, M. J. & Herrick, R. Evaluating fungal populations by genera/species on wide body commercial passenger aircraft and in airport terminals. Annals Occup. Hyg. 51, 281–291, DOI: https://doi.org/10.1093/annhyg/mem002 (2007).

14. Schaumburg, F., Köck, R., Leendertz, F. H. & Becker, K. Airport door handles and the global spread of antimicrobial-resistant bacteria: a cross sectional study. Clin. Microbiol. Infect. 22, 1010–1011, DOI: https://doi.org/10.1016/j.cmi.2016.09.010 (2016).

15. Verelst, F., Willem, L. & Beutels, P. Behavioural change models for infectious disease transmission: a systematic review (2010–2015). J. Royal Soc. Interface 13, 20160820, DOI: http://dx.doi.org/10.1098/rsif.2016.0820 (2016).

16. Poletti, P., Ajelli, M. & Merler, S. The effect of risk perception on the 2009 H1N1 pandemic influenza dynamics. PLOS ONE 6, 1–7, DOI: https://doi.org/10.1371/journal.pone.0016460 (2011).

17. Greenwood, B. The contribution of vaccination to global health: past, present and future. Philos Trans R Soc Lond B Biol Sci 369, 20130433, DOI: 10.1098/rstb.2013.0433 (2014).

18. Yang, Y. et al. The transmissibility and control of pandemic influenza A (H1N1) virus. Science 326, 729–733, DOI: 10.1126/science.1177373 (2009). http://science.sciencemag.org/content/326/5953/729.full.pdf.

19. Yokum, D., Lauffenburger, J. C., Ghazinouri, R. & Choudhry, N. K. Letters designed with behavioural science increase influenza vaccination in medicare beneficiaries. Nat. Hum. Behav. 2, 743–749, DOI: 10.1038/s41562-018-0432-2 (2018).

20. Patel, M. S. Nudges for influenza vaccination. Nat. Hum. Behav. 2, 720–721, DOI: 10.1038/s41562-018-0445-x (2018).

21. Huizer, Y. L., Swaan, C. M., Leitmeyer, K. C. & Timen, A. Usefulness and applicability of infectious disease control measures in air travel: a review. Travel. Medicine Infect. Dis. 13, 19–30, DOI: https://doi.org/10.1016/j.tmaid.2014.11.008 (2015).

22. Nicolaides, C., Cueto-Felgueroso, L. & Juanes, R. The price of anarchy in mobility-driven contagion dynamics. J. The Royal Soc. Interface 10, DOI: 10.1098/rsif.2013.0495 (2013). http://rsif.royalsocietypublishing.org/content/10/87/20130495.full.pdf.

23. Aiello, A. E. & Larson, E. L. What is the evidence for a causal link between hygiene and infections? The Lancet Infect. diseases 2, 103–110, DOI: https://doi.org/10.1016/S1473-3099(02)00184-6 (2002).

24. Wong, V. W. Y., Cowling, B. J. & Aiello, A. E. Hand hygiene and risk of influenza virus infections in the community: a systematic review and meta-analysis. Epidemiol. Infect. 142, 922–932, DOI: 10.1017/S095026881400003X (2014).

25. Aiello, A. E., Coulborn, R. M., Perez, V. & Larson, E. L. Effect of hand hygiene on infectious disease risk in the community setting: A meta-analysis. Am. J. Public Heal. 98, 1372–1381, DOI: https://doi.org/10.2105/AJPH.2007.124610 (2008).

26. Rabie, T. & Curtis, V. Handwashing and risk of respiratory infections: a quantitative systematic review. Trop. Medicine & Int. Heal. 11, 258–267, DOI: 10.1111/j.1365-3156.2006.01568.x (2006). https://onlinelibrary.wiley.com/doi/pdf/10.1111/j.1365-3156.2006.01568.x.

27. Null, C. et al. Effects of water quality, sanitation, handwashing, and nutritional interventions on diarrhoea and child growth in rural Kenya: a cluster-randomised controlled trial. The Lancet Glob. Heal. 6, e316–e329, DOI: 10.1016/S2214-109X(18) 30005-6 (2018).

28. World Health Organisation. Mitigating the impact of the new influenza a(h1n1): options for public health measures. http://www.wpro.who.int/emerging_diseases/documents/Mitigating10June.pdf. Online; accessed October 2018.

29. Burton, M. et al. The effect of handwashing with water or soap on bacterial contamination of hands. Int J Environ Res Public Heal. 8, 97–104, DOI: 10.3390/ijerph8010097 (2011).

30. Colizza, V., Pastor-Satorras, R. & Vespignani, A. Reaction-diffusion processes and metapopulation models in heterogeneous networks. Nat. Phys. 3, 276–282, DOI: https://doi.org/10.1038/nphys560 (2007).

31. Brockmann, D. & Helbing, D. The hidden geometry of complex, network-driven contagion phenomena. Science 342, 1337–1342, DOI: 10.1126/science.1245200 (2013).

32. Nicolaides, C., Cueto-Felgueroso, L., González, M. C. & Juanes, R. A metric of influential spreading during contagion dynamics through the air transportation network. PLoS ONE 7, e40961, DOI: https://doi.org/10.1371/journal.pone.0040961 (2012).

33. González, M. C., Hidalgo, C. A. & Barabási, A.-L. Understanding individual human mobility patterns. Nature 453, 779–782, DOI: https://doi.org/10.1038/nature06958 (2008).

34. Barnhart, C., Fearing, D. & Vaze, V. Modeling passenger travel and delays in the national air transportation system. Oper. Res. 62, 580–601, DOI: https://doi.org/10.1287/opre.2014.1268 (2014).

35. Vespignani, A. Modelling dynamical processes in complex socio-technical systems. Nat. Phys. 8, 32–39, DOI: http://dx.doi.org/10.1038/nphys2160 (2011).

36. Brauer, F. Mathematical epidemiology: Past, present, and future. Infect. Dis. Model. 2, 113–127, DOI: https://doi.org/10.1016/j.idm.2017.02.001 (2017).

37. Lee, M.-S., Hong, S. J. & Kim, Y.-T. Handwashing with soap and national handwashing projects in Korea: focus on the National Handwashing Survey, 2006-2014. Epidemiol. Heal. 37, e2015039, DOI: https://doi.org/10.4178/epih/e2015039 (2015).

38. American Society for Microbiology. Another US airport travel hazard - dirty hands (2003). Retrieved from https://www.eurekalert.org/pub_releases/2003-09/asfm-aua091103.php; accessed October 2018.

39. Centers for Disease Control and Prevention. When & how to wash your hands. https://www.cdc.gov/handwashing/when-how-handwashing.html (2016). Online; accessed October 2018.

40. Borchgrevink, C. P., Cha, J. & Kim, S. Hand washing practices in a college town environment. J. Environ. Heal. 75, 18–24 (2013).

41. Merk, H., Kühlmann-Berenzon, S., Linde, A. & Nyrén, O. Associations of hand-washing frequency with incidence of acute respiratory tract infection and influenza-like illness in adults: a population-based study in sweden. BMC Infect. Dis. 14, 509, DOI: 10.1186/1471-2334-14-509 (2014).

42. Rothman, K. J. Epidemiology: An Introduction (Oxford University Press, New York, USA, 2012).

43. Smith, R. D. Responding to global infectious disease outbreaks: Lessons from sars on the role of risk perception, communication and management. Soc. Sci. & Medicine 63, 3113–3123, DOI: https://doi.org/10.1016/j.socscimed.2006.08.4 (2006).

44. Fan, V. Y., Jamison, D. T. & Summers, L. H. Pandemic risk: how large are the expected losses? Bull. World Heal. Organ. 96, 129–134, DOI: http://dx.doi.org/10.2471/BLT.17.199588 (2018).

45. Chung, L. H. Impact of pandemic control over airport economics: Reconciling public health with airport business through a streamlined approach in pandemic control. J. Air Transp. Manag. 44–45, 42–53, DOI: https://doi.org/10.1016/j.jairtraman.2015.02.003 (2015).

46. Ferguson, N. M. et al. Strategies for mitigating an influenza pandemic. Nature 442, 448–452, DOI: http://dx.doi.org/10.1038/nature04795 (2006).

47. Airbus. A330neo Family: Powering into the future. https://www.airbus.com/content/dam/corporate-topics/publications/backgrounders/Backgrounder-Airbus-Commercial-Aircraft-A330neo-E.pdf. (2018). Online; accessed October 2018.

48. Boeing. The airplane bathroom that cleans itself. https://www.boeing.com/features/2016/03/self-clean-lavatory-03-16.page (2016). Online; accessed October 2018.

49. Bormann, R., Weisshardt, F., Arbeiter, G. & Fischer, J. Autonomous dirt detection for cleaning in office environments. In 2013 IEEE International Conference on Robotics and Automation, 1260–1267, DOI: 10.1109/ICRA.2013.6630733 (2013).

50. Smixin. Automatic and connected hand washing system from Smixin. http://www.smixin.com/automatic-and-connected-hand-wash-system-from-smixin/ (2017). Online; accessed October 2018.

51. Aral, S. & Nicolaides, C. Exercise contagion in a global social network. Nat. Commun. 8, 14753, DOI: http://dx.doi.org/10.1038/ncomms14753 (2017).

52. Lim, J. & Meer, J. How do peers influence BMI? evidence from randomly assigned classrooms in South Korea. Soc. Sci. & Medicine 197, 17–23, DOI: https://doi.org/10.1016/j.socscimed.2017.11.032 (2018).

53. Grover, E. et al. Social influence on handwashing with soap: Results from a cluster randomized controlled trial in Bangladesh. The Am. J. Trop. Medicine Hyg. 99, 934–936, DOI: https://doi.org/10.4269/ajtmh.17-0903 (2018).

